# Opposing effects of floral visitors and soil conditions on the determinants of competitive outcomes maintain species diversity in heterogeneous landscapes

**DOI:** 10.1101/170423

**Authors:** Jose B. Lanuza, Ignasi Bartomeus, Oscar Godoy

## Abstract

Theory argues that both soil conditions and aboveground trophic interactions are equally important for determining plant species diversity. However, it remains unexplored how they modify the niche differences that stabilise species coexistence and the average fitness differences driving competitive dominance. We conducted a field study in Mediterranean annual grasslands to parameterise population models of six competing plant species. Spatially explicit floral visitor assemblages and soil salinity variation were characterized for each species. Both floral visitors and soil salinity modified species population dynamics via direct changes in seed production and indirect changes in competitive responses. Although the magnitude and sign of these changes were species specific, floral visitors promoted coexistence at neighbourhood scales while soil salinity did so over larger scales by changing the superior competitor's identity. Our results show how below and aboveground interactions maintain diversity in heterogeneous landscapes through their opposing effects on the determinants of competitive outcomes.

## Introduction

Understanding the mechanisms that maintain species diversity is a central aim in ecology. Although species interact with the environment and with many other species in complex ways, ecologists have traditionally assumed that the importance of biotic and abiotic factors in promoting species diversity is highly asymmetrical. Competition driven by soil conditions is commonly considered to be the primary driver of plant coexistence, and therefore it has been extensively explored (Raynaud & Leadley 2004; Tilman 2006; Craine & Dybzinski 2013; Hendriks *et al.* 2015). For instance, differences in the species’ ability to drawdown limiting resources such as nitrogen and phosphorous is a classic textbook example illustrating the importance of partitioning soil resources for maintaining species diversity (Tilman 1994). On top of this, multitrophic interactions such as those occurring between plant and pollinators, pathogens, or mycorrhizae (Fitter 1977; Bastolla *et al.* 2009; Bagchi *et al.* 2014; Parker *et al.* 2015; Bennett *et al.* 2017), are thought to play a secondary role in structuring plant communities.

However, recent work has challenged this view. Chesson & Kuang (2008) presented clear evidence that there is no theoretical support that the relative importance of competition driven by soil conditions versus other kinds of multitrophic interactions is asymmetrical. In fact, they argue that these two types of interactions are equally able to either limit or promote diversity. Competition mediated by other trophic levels has been largely studied under the concept of *apparent competition* (Holt 1977), which specifically describes how species within a trophic level (e.g. plants) can produce indirect competitive effects on others via shared enemies (e.g. herbivores, predators). These indirect effects can be of equal magnitude to that of direct effects resulting from resource competition. The utility of this concept has also been extended to apparent negative and positive interactions mediated by other organisms, such as shared mutualisms (Morris *et al.* 2004; Carvalheiro *et al.* 2014).

Plant interactions with floral visitors, including pollinators and pollen thieves, are key trophic interactions with clear potential to strongly influence plant coexistence (Morris *et al.* 2010; Ollerton *et al.* 2011). Many theoretical studies have suggested several mechanisms by which floral visitors can promote plant diversity (Bastolla *et al.* 2009; Benadi *et al.* 2013; Pauw 2013). For instance, Fontaine *et al.* (2005) shows experimentally that plant richness correlates positively with greater functional diversity of floral visitors. Yet the effect of floral visitors on plant coexistence remains poorly understood, as their effects on plant population dynamics have not been related to the determinants of competitive outcomes.

Similar to previous work focused on multitrophic antagonistic interactions (mainly predators and pathogens) (Chesson & Kuang 2008; Kuang & Chesson 2010; Stump & Chesson 2017), we can obtain progress by framing our research within recent advances of coexistence theory (Chesson 2000). According to Chesson’s framework, floral visitors can promote the *stabilising niche differences* that favour plant coexistence, which occur when intraspecific competitive interactions exceed interspecific competition, and the *average fitness differences* that favour competitive exclusion, and determine the competitive winner in the absence of niche differences. Ecologists have paid much more attention to the floral visitors’ effects on average fitness differences (Herrera 2000; Thompson 2001; Waites & Ågren 2004), than their effect on stabilising niche differences (Pauw 2013). However, as most plants depend on pollinators to maximize their reproductive success (Ollerton *et al.* 2011), it is most likely that floral visitors’ characteristics may promote both plant niche and fitness differences simultaneously. Therefore, the effect of floral visitors on determining plant coexistence is only predictable with a mechanistic understanding of how different assemblages modify niche and fitness differences. Coexistence can be achieved by several pathways, either by equalising fitness differences, by promoting niche differences, or by a combination of both.

Relating field data to these theoretical advances is challenging but we can take advantages of systems relatively easy to observe for which recent work has described how niche and fitness differences influences species’ population dynamics, such Mediterranean annual grasslands (Godoy & Levine 2014). Floral visitor assembleges in these environments are particularly interesting because they are composed of an array of insects including solitary bees, hover flies, beetles, and butterflies, which can produce contrasting effects on plant fitness. The fitness of many plant species relies on those floral visitors that are truly pollinators (Pauw 2013), and other insects can have a negative effect on plant fitness by robbing their nectar, eating pollen or by damaging the flower (Morris *et al.* 2003). Mediterranean annual grasslands also often inhabit in salty soils. Variation in saline soil conditions is negatively correlated with soil fertility (Olff & Ritchie 1998; Hu & Schmidhalter 2005) and similar to floral visitors salt concentration on soils can have contrasting effects on plant fitness. While glycophytes will struggle to grow under saline conditions (Flowers & Yeo 1986) halophytes are expected to show greater fitness and competitive advantages (Flowers & Colmer 2008).

Here, we considered three interaction levels to test how the belowground environmental conditions (i.e. soil conditions) and the aboveground trophic interactions (i.e. floral visitors) influence coexistence of the middle level (i.e. plant species). We specifically focus on three questions: (1) How soil salinity and floral visitors modify species’ population dynamics via direct changes in per capita seed production and indirect changes in species’ responses to competitive interactions? (2) Do these direct and indirect effects modify niche and fitness differences between plant species?, and finally, (3) Are these modifications of the determinants of competitive outcomes limiting or promoting diversity?

We answered these three questions by first parameterizing a general plant competition model from which the stabilising niche differences and average fitness differences were quantified. To parameterise these models of pairwise competition between six annual European grassland species, we quantified their vital rates (i.e. germination, fecundity, seed survival) and competition coefficients in field plots relating seed production of focal individuals to a density gradient of numerous different competitors. We then assessed how seasonal and spatial variation in the number of floral visits and in soil salinity changes species fecundity and their responses to competition (Question 1). Once, the model was parameterised, we estimated niche and fitness differences when soil salinity and floral visitors were present or absent (Question 2), and compared how strong niche differences offset fitness differences between scenarios (Question 3). Our work is novel in quantifying the effects that distinct environmental conditions and trophic interactions have on modifying niche and fitness differences, and showing under field conditions that they maintain diversity in heterogeneous landscapes through their opposing effects on the determinants of competitive outcomes.

## Methods

### Study system

Our study was conducted in Caracoles Ranch (2,680 ha), an annual grassland system located in Doñana NP, southwest Spain (37°04’01.5“N 6°19’16.2”W). The climate is Mediterranean with mild winters and average 50-y annual rainfall of 550-570 mm with high interannual oscillations (Muñoz-Reinoso & García Novo 2000). Soils are sodic saline (electric conductivity > 4dS/m and pH < 8.5) and annual vegetation dominates the grassland with no perennial species present. The study site has a subtle micro topographic gradient (slope 0.16%) enough to create vernal pools at lower parts from winter (November-January) to spring (March-May) while upper parts do not get flooded except in exceptionally wet years. A strong salinity-humidity gradient is structured along this topographic gradient. Additionally, salt can reach upper parts of the soil by capillarity during the rainfall period resulting overall in heterogeneous soil salinity patterns at the local and at the landscape scale (Appendix S1). This salinity gradient is strongly correlated with soil nutrient availability at our study location, and more saline soil conditions were less fertile especially in phosphorous content (Clemente *et al.* 2004).

We initially focused on 16 annual plants that were common at the study site. These species represent a broad range of taxonomic families, plant morphology and flowering phenology co-occurring at the scale of the entire study system. All species were considered for estimating competitive interactions, but we observed enough visits of insects to the flowers of six species. Hence, we further focus on this particular set of species to compare the effect of soil salinity and floral visitors on niche and fitness differences (Table 1).

**Table 1.**
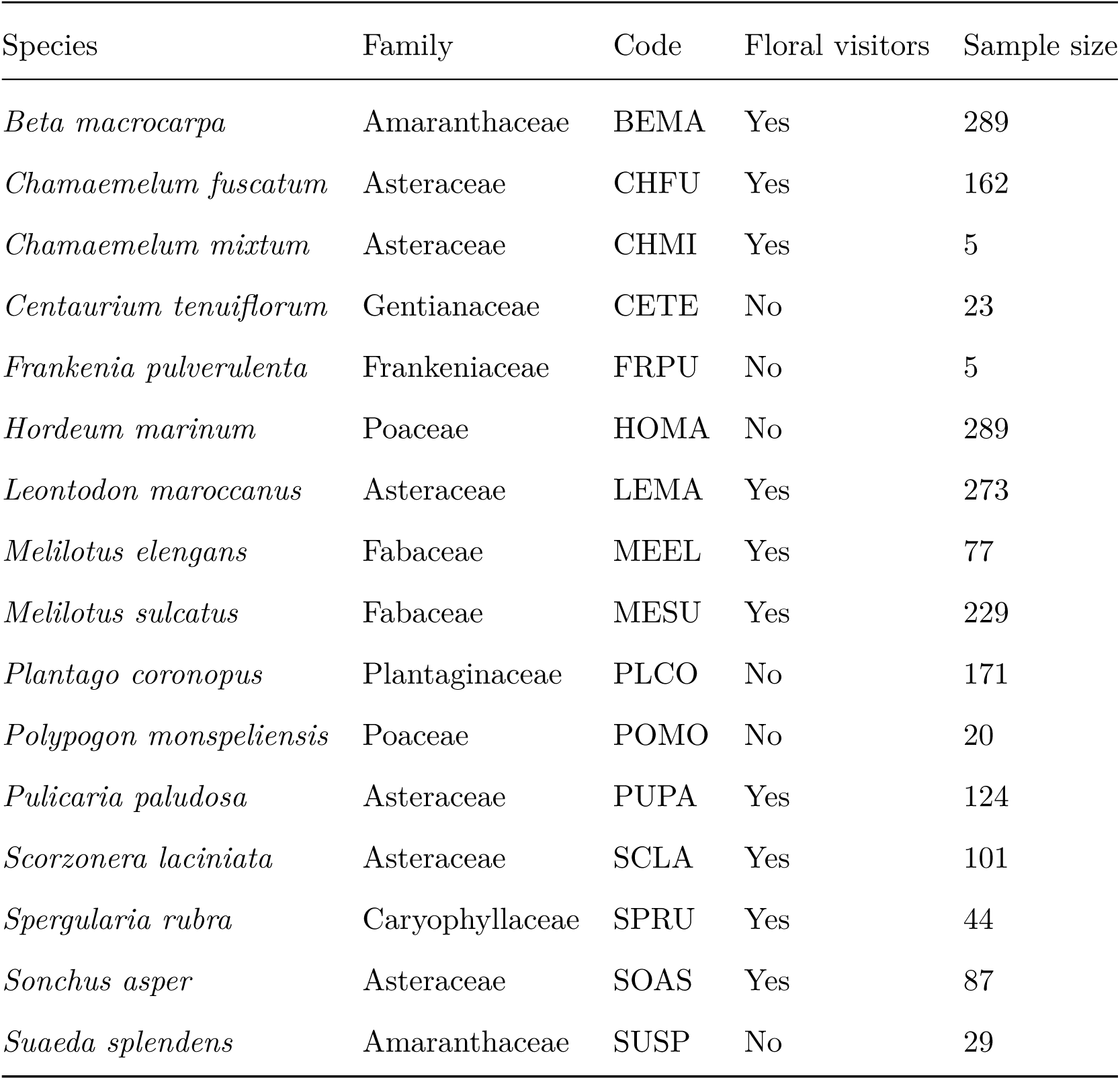
List of species observed in Caracoles Ranch. Code and taxonomic family of each species is provided. Sample size represents the total number of individuals sampled for each focal species, and it is correlated with their natural abundance at the study site.

### Modeling approach to quantify the niche and fitness differences between species pairs

Our observational study was designed to field-parameterise a mathematical model describing annual plant population dynamics (Levine & HilleRisLambers 2009). This model allows quantifying stabilising niche differences and average fitness differences between species within a trophic level (Godoy & Levine 2014). Importantly, there have not been previous attempts to quantify how soil condition or multitrophic interactions change the strength of niche and fitness differences between species within a single trophic level, and here we show how these effects can be incorporated into this model. The model is described as follows:

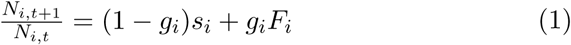

where 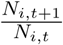 is the per capita population rate, and *N*_*i*,*t*_ is the number of seeds of species *i* in the soil prior to germination in winter of year *t*. The germination rate of species *i*, *g_i_*, can be viewed as a weighting term for an average of two different growth rates: the annual survival of ungerminated seed in the soil (*s_i_*), and the viable seeds produced per germinated individual (*F_i_*). In past work, *F_i_*, was expanded into a function describing how the average fecundity of each germinated seed that becomes an adult (i.e. per germinant fecundity) declines with the density of competing number individuals in the system (Godoy & Levine 2014). Now, we slightly modify this function to include the additional effect of floral visitors and soil conditions on the per germinant fecundity as follows:

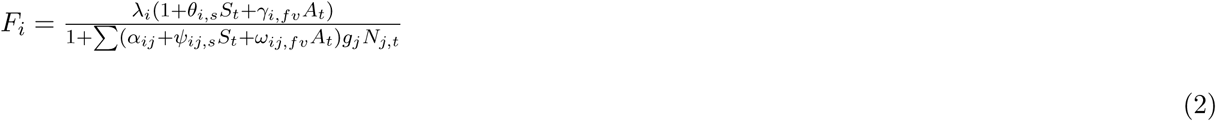

where *θ*_*i*,*s*_ and *γ*_*i*,*fv*_ control the effect of soil salinity (*S_t_*) and floral visitors (*A_t_*) respectively on the per germinant fecundity of species *i* in the absence of competition (λ_*i*_). In addition, λ_*i*_ is modified by the germinated densities of other species including its own (*g_j_N*_*j*,*t*_). To describe the per capita effect that species j is mediating on species i, we multiplied these germinated densities by a sum of three interaction coefficients (*α_ij_*+*ψ*_*ij*,*s*_+*ω*_*ij*,*fv*_), which describes the additional direct effect of soil salinity and the apparent effect of floral visitors on the competitive interactions between species. Notice that we considered only explicitly in our study the e**ff**ect that soil salinity and floral visitors have on species’ fecundity (*F_i_*), but the model could be easily extended to include the effect of these two factors on the other two vital rates, germination (*g_i_*) and seed soil survival (*s_i_*).

With the direct and apparent dynamics of competition described by this population model, we followed the approach of Chesson (2012) to determine fitness and niche differences between species pairs. Our procedure here parallels previous work described in Godoy & Levine (2014), and allows us to define stabilising niche differences and fitness differences with and without considering the effect of floral visitors and soil salinity on plant coexistence. For the model described by eqns (1) and (2), we define niche overlap (*ρ*) as follows:

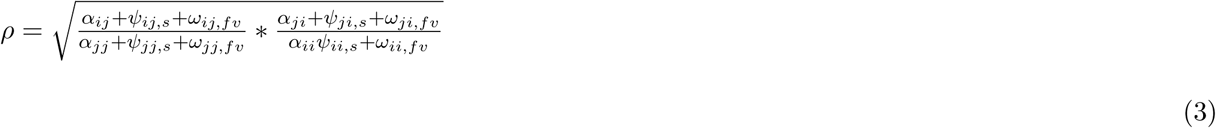

If multitrophic interactions are absent (i.e. *ψ*_*ij*,*s*_=0, *ω*_*ij*,*fv*_=0), then *ρ* collapses into an equation that reflects the average degree to which species limit individuals of their own species relative to heterospecific competitors based on their interaction coefficients (*α*’s) (Godoy & Levine 2014). Conversely, if multitrophic interactions are present *ψ* and *ω* are the terms controlling changes in average niche differences between a pair of species. For example, two species with different set of floral visitors could increase niche overlap by having positive apparent competitive effects of each species on the other (i.e *ω*_*ij*,*fv*_ >0). With (*ρ*) defining niche overlap between a pair of species, their stabilising niche difference is expressed as 1-*ρ*.

As an opposing force to stabilising niche differences, average fitness differences drive competitive dominance, and in the absence of niche differences, determine the competitive superior between a pair of species. Addressing the modifications done in the annual population model described by eqns (1) and (2) to include the effect of floral visitors and soil conditions, we define average fitness differences between the competitors 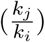 as:

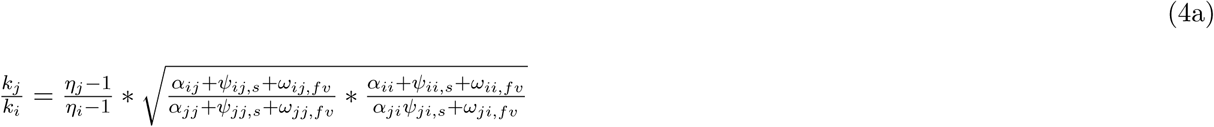

and

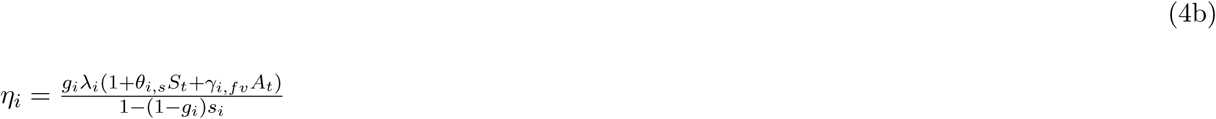

When the ratio 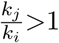 this indicates that species *j* has a fitness advantage over species *i.* Both soil salinity and floral visitors can be seen as equalising mechanisms promoting coexistence because they can reduce fitness differences between a species pair by two contrasted pathways. They can modify the ‘demographic ratio’ 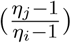 which describes the degree to which species *j* produces more seeds (*g_j_*λ_*j*_(1 + *θ*_*j*,*s*_*S_t_* + *γ*_*j*,*fv*_*A_t_*)) per seed loss due to death or germination (1-(1-*g_j_*)*s_j_*) than species *i*, and they can also modify the ‘competitive response ratio’ 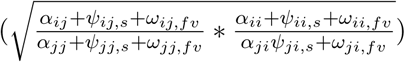 which describes the degree to which species *j* is less sensitive to competition than species *i* (eqn (4)). Notice that these modifications can produce the opposing effect and promote species’ competitive dominance by a combination of high demographic rates and low sensitivity to competition.

Competitors can coexist when niche differences overcome fitness differences, allowing both species to invade (i.e. increase its populations) when rare (Chesson 2012). This condition for mutual invasibility is statisfied when:

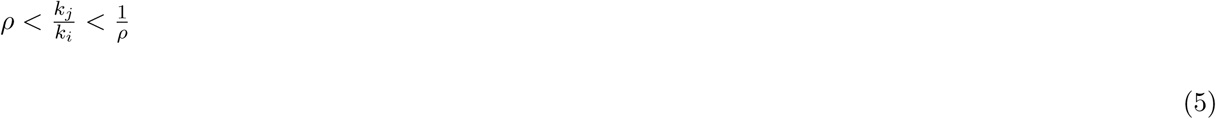

Therefore, coexistence can occur when little niche overlap (i.e. large niche differences) overcomes large fitness differences, or when niche differences among similar species are still large enough to stabilise the interaction between competitors with similar fitness. We used this condition to evaluate how strongly floral visitors and soil salinity increase or decrease the likelihood of coexistence between competitors. For doing so, we computed how much observed niche differences exceed or fails to promote coexistence according to the expected niche differences needed to overcome observed fitness differences between a species pair.

### Field observations used to parameterise the model

In September 2015, we established nine plots of 8.5m × 8.5m along a 1km × 200m area following a topographic gradient. Three of these nine plots where located in the upper part of the topographic gradient, three at the middle, and the last three at the lower part. Average distance between these three locations was 300m and average distance between plots within each location was 15m. In addition, each plot was divided in 36 subplots of 1m × 1m with aisles of 0.5m in between to allow access to subplots where measurements were taken (Appendix S2).

This spatial design was established to parameterise models of pairwise competition between the six focal species finally considered with estimates of species’ germination fractions (*g_i_*), seed survival in the soil (*s_i_*), and the effect of soil salinity (*S_t_*) and floral visitors (*A_t_*) on the per germinant fecundities in the absence of neighbours (λ_*i*_) and on all pairwise interaction coefficients (*α_ij_*). Specifically, the core of the observations involved measuring per germinant viable seed production as a function of the number and identity of neighbours within a radius of 7.5cm including individuals of the same species (see analyses below). We measured one individual per subplot for widespread species and several individuals per subplot when species were rare (max. 324 individuals/species). To additionally incorporate the effect of soil salinity, we measured from November 2015 to June 2016 soil humidity (%) and soil salinity (dS/m) bimonthly at the subplot center with a TDR (Time Domain Reflectometer) incorporating a 5cm probe specially designed and calibrated for these sodic saline soils (EasyTest, Poland). We summarised the amount of soil salinity experienced by each germinant, which was highly correlated with soil moisture (r=0.77), as the sum over their lifetime of the soil salinity measured at the subplot scale.

Moreover, floral visitors were measured during the complete phenological period of all species (from January to June 2016). We surveyed weekly the number of floral visitors for all species within each subplot. Visits were only considered when the floral visitor touched the reproductive organs of the plant. All subplots within a plot were simultaneously surveyed during 30 minutes each week. Unknown floral visitors were collected with a hand net and the time spent collecting the insects was discounted from the total observation time in order to equalise the amount of time used per plot. Plot survey was randomized between weeks to avoid sampling effects. Overall, this procedure rendered approximately 90 hours of overall floral visitors sampling. Floral visitors to each species and subplot were grouped in four main morphology groups (bees, beettles, butterflies, and flies). We summarised the number of floral visits by insects to each germinant as the total sum of visits at the subplot scale.

Finally, we quantified the germination of viable seeds (*g_i_*) by counting the number of germinants in 18 quadrats of 1m x 1m placed close to the plots (2 quadrats per plot) from seeds collected the previous year and sown on the ground prior to the first major storm event after summer (September 2015). Similarly, we quantified seed bank survival (*s_i_*) with the same seed material by burying seeds from September 2015 to September 2016 following the methods of (Godoy & Levine 2014).

### Analysis

To fit the model to the empirical observations, we used maximum likelihood methods (optim R function, method=“L-BFGS-B”). We fit changes in λ_*i*_ and *α_ij_* (both bounded to be positive) as a function of soil salinity (*S_t_*) and floral visitors (*A_t_*). Soil salinity (*θ*_*i*,*s*_, *ψ*_*ij*,*s*_) and floral visitors (*γ*_*i*,*fv*_, *ω*_*ij*,*fv*_) parameters were not bounded to any specific range as we hypothesized that they can have both positive and negative effects on per germinant fecundities. What we do not know, however, is whether their effects are specific to each pairwise interactions or are common to all species interactions. To test for this possibility, we use an AIC (Akaike Information Criteria) model selection approach to distinguish which of two following models were the best fit for our observations for each target species *i*. The first model (model 1) assumes that competitive interactions between species are pairwise specific but the effects of salt and floral visitors on competitive interactions are common across species.

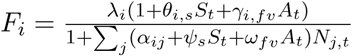

The second model (model 2) assumes that competitive interactions between species are pairwise specific, as are the effects of salt and floral visitors on competitive interactions.

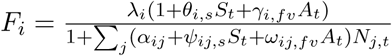

It is also likely that soil salinity and floral visitors may not be affecting the competitive dynamics between species pairs. Therefore, we evaluated an additional model (model 3) that did not account for these abiotic and biotic factors.

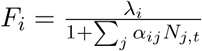

For all three models, individuals of the 10 species surveyed apart from our six focal species were grouped together and their competitive effect on the six focal species was summarized as a single parameter. The average viable seed production per species (*F_i_*) was estimated by counting the number of fruits produced by 20 independent germinants collected across plots, counting the number of seeds produced per fruit, and correcting for its viability. *A_t_* and *S_t_* represent the total number of floral visits and the accumulated soil salinity experienced by each germinant at the subplot scale over their lifetime. Estimates of mean and standard error for each parameter of the best model selected by AIC are included in Appendix S3. All analyses were conducted in R (version 3.3.1) (R Core Team 2016).

## Results

The six focal species experienced a great variation in soil salinity and the type and number of floral visitors. Along the salinity gradient, *Beta macrocarpa* and *Pulicaria paludosa* grew mainly in high soil salinity concentrations, in contrast, *Melilotus elegans* and *Leontodon maroccanus* grew in relatively low saline soils, while *Chamaemelum fuscatum* and *Melilotus sulcatus* showed a more tolerant behaviour growing in a wider range of salt concentrations (Fig. 1). Number of floral visits by insects also varied greatly among plant species. Overall, the main groups of floral visitors in our system were flies (581 visits) and beetles (496 visits), followed by bees (161 visits) and butterflies (87 visits). The three Asteraceae species were the most visited species. Among them, *L. maroccanus* received 636 visits followed by *C. fuscatum* 293 visits, and *P. paludosa* 291 visits. The rest of the species *B. macrocarpa* (64), *M. sulcatus* (35), and *M. elegans* (6) had in comparison a much lower number of visits. Moreover, species also showed variation in the assemblage of floral visitors. Of the three plant species with higher number of visits, flies were the most abundant insects visiting *C. fuscatum*, and *P. paludosa* while beetles did so for *L. maroccanus* (Fig. 1).

**FIGURE 1.**
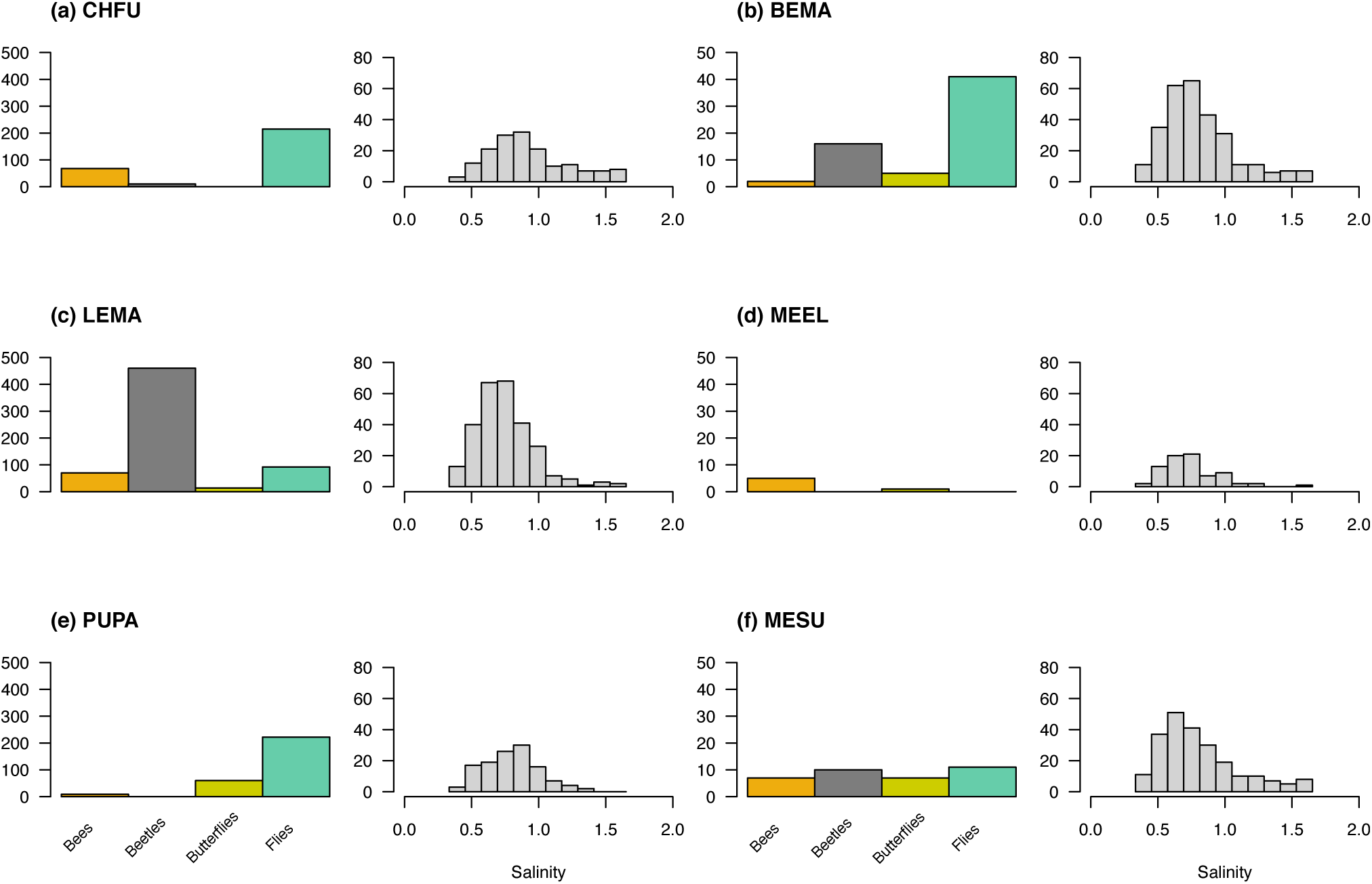
For the six focal species this shows: total number of visits of the four groups of floral visitors (bees, beetles, butterflies and flies) (left panel) and species abundance along the salinity gradient (right panel). The amount of salinity experience during the life span of each species was measured as the sum of the electric conductivity in Ds/m measured bi-monthly. Note that the three Asteraceae species (a) CHFU, (c) LEMA and (e) PUPA had an order of magnitude more floral visits than the non-Asteraceae species (b) BEMA, (d) MEEL and (f) MESU. See Table 1 for species code.

The wide variation in the number of floral visits and soil salinity concentrations observed in our experiment modified the seed production in the absence of neighbours (λ_*i*_) and the strength of the species’ responses to competitive interactions (*α_ij_*) of the three Asteraceae species (model 2, lowest AIC values) (Appendix S4). Interestingly, the sign of the floral visitors’ effects on λ_*i*_ and *α_ij_* varied among these species. While higher number of visits to *C. fuscatum* increased its potential fecundity and reduced the negative effect of both intra and interspecific competition on seed production, the opposite pattern was observed for *L. maroccanus* and *P. paludosa* (Fig. 2 and 3). Soil salinity, in contrast, had a similar effect across species increasing seed production in the absence of neighbours and promoting weaker competitive interactions. For the other three non Asteraceae species, AIC values did not help to distinguish unambiguously whether soil salinity and floral visitors had a common effect on λ_*i*_ and *α_ij_* (i.e. differences in AIC between model one and three were lower than 10). In neither case, model selection did support the view that the effect of floral visitors and soil salinity on the species’ responses to competition was pairwise specific (i.e. model two showed consistently higher AIC values) (Appendix S4).

**FIGURE 2.**
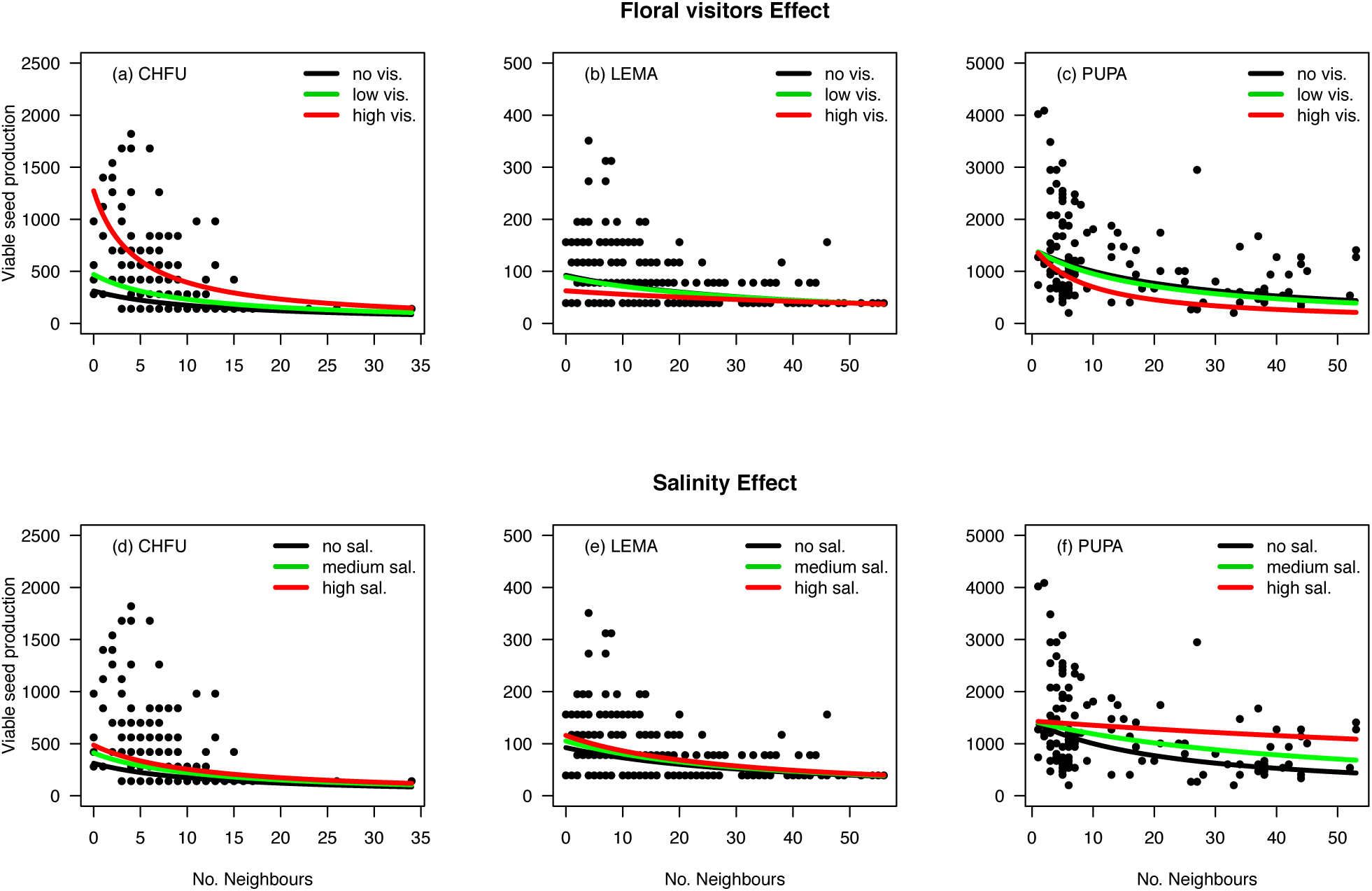
Relationship between per capita seed production as a function of the number of neighbours within a 7.5 cm radius area for the six studied plant species. The negative exponential regressions are represented with the parameters estimated from the maximum likelihood approach and median conditions of soil salinity and frequency of floral visitors experienced by each focal species. Parameters estimates correspond to the AIC best-supported model, which were model 1 for (a) CHFU, (c) LEMA and (e) PUPA and model 3 for (b) BEMA, (d) MEEL and (f) MESU. Intraspecific competitive effects are represented with a solid black line whereas interspecific effects are represented with a coloured dashed line.

**FIGURE 3.**
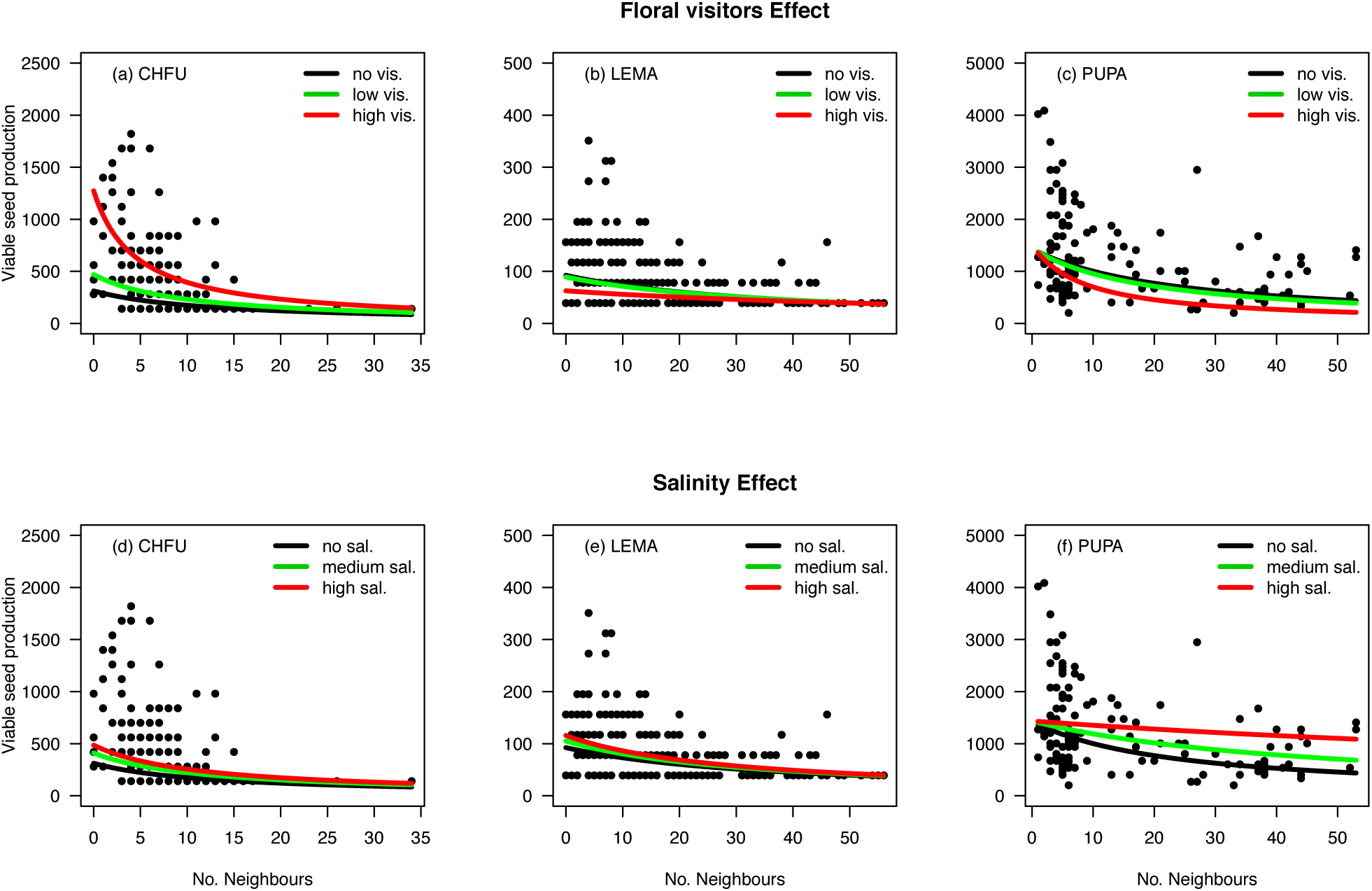
Relationship between per capita seed production as a function of the number of neighbours according to three different conditions of floral visitors and soil salinity. Here is shown the three focal species (CHFU, LEMA, and PUPA) for which these abiotic and biotic factors had a significant common effect on species’ fecundity (model 1). Upper panel contains floral visitors effects with black curves representing no floral visitation, green curves representing one or two visits, and red curves representing percentile 95 of floral visits (which ranges from 6 visits in *C. fuscatum* to 9 visits in *L. maroccanus* and *P. paludosa*). Lower panel contains soil salinity effects with black curves considering no salt in the soil, and green and red curves representing percentiles 50 and 95 respectively of the soil salinity sum over focal species life span.

Soil salinity and floral visitors exerted positive, negative or no effect on plant fitness, yet they modified the determinants of competitive outcomes in opposite and specific directions (Fig. 4). While floral visitors tend to maintain stable coexistence (5 out of 15 species pairs) or to promote coexistence by equalising fitness differences (5 out of 15 species pairs moved closer to the coexistence region), soil salinity tend to promote competitive exclusion (4 species pairs moved out of the coexistence region) and increase competitive asymmetries between species pairs. As a result, floral visitors reduced on average the niche differences needed for coexistence (estimated from the mutual invasibility, eqn (5)) across species pairs (paired t-test, t = 2.15, P = 0.049), while soil salinity increased significantly the niche differences needed for coexistence (paired t-test, t = 5.51, P < 0.001). Although soil salinity reduced the likelihood of species coexistence at neighbourhood scales for all except one species pair, this abiotic factor also determined changes in the identity of competitive winners (6 out of 15 species pairs), which suggest that soil salinity can promote species coexistence over larger scales by turnover of the dominant competitor (Fig. 4).

**FIGURE 4.**
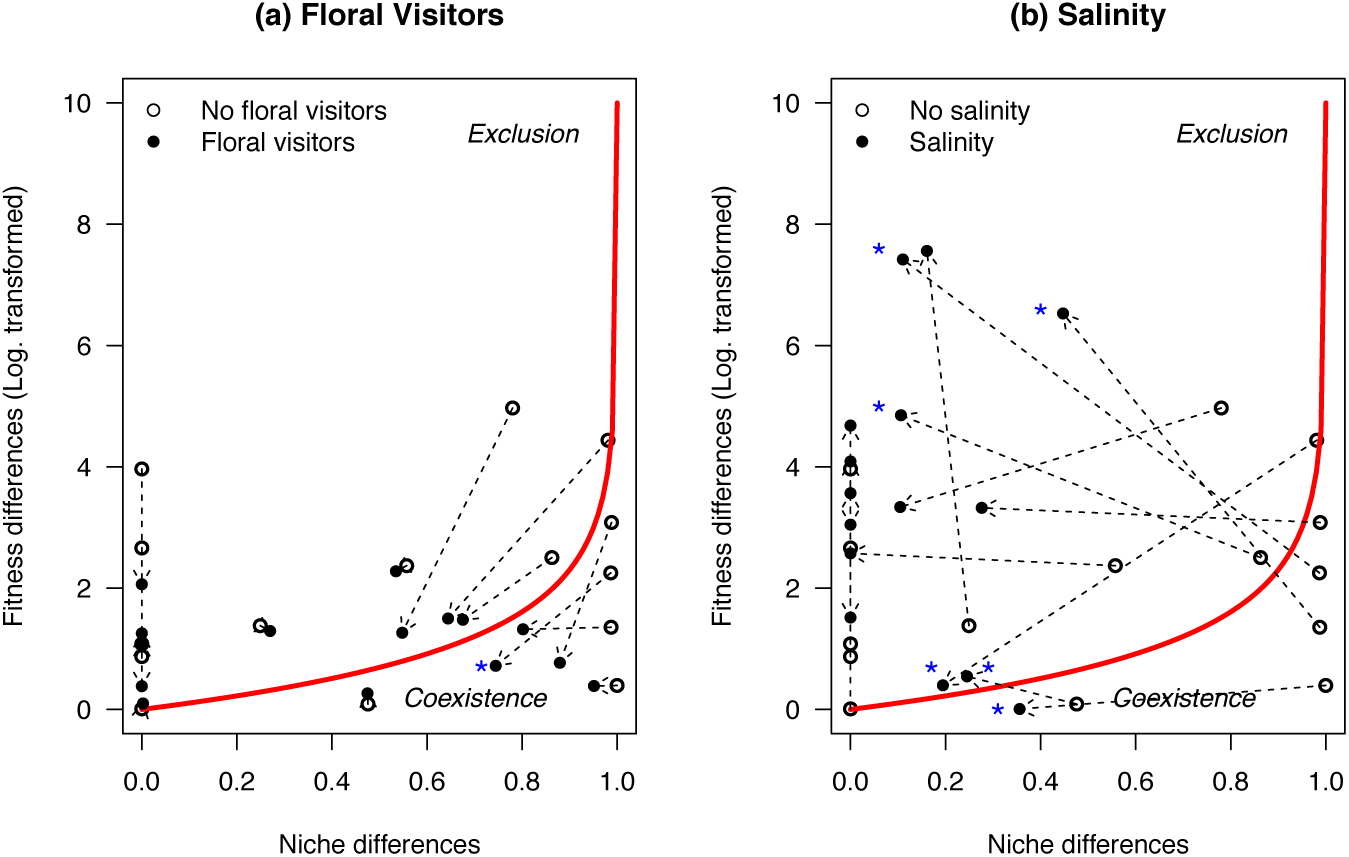
Average fitness and stabilising niche differences for each pair of species (denoted by a single point). Black solid points correspond to the situation of considering the effect of floral visitors (a) or soil salinity (b) on the determinants of competition outcomes, and white open points correspond to the situation of not considering these effects. Dashed arrows connect both situations within species pairs. The red curve separates the exclusion region from the region where the condition for coexistence is met (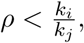, where species j is the fitness superior). Five species pairs fall in the coexistence region without considering the effect of floral visitors and soil salinity. When considering the effect of floral visitors, these five coexisting pairs were remained within the coexistence region and another five moved closer to the coexistence region. In contrast, soil salinity moved out from the coexistence region four of the five coexisting species pairs away from the coexistence region and increased the degree of competitive asymmetry between eight species pairs more. Nevertheless, soil salinity changed the identity of the competitive superior in six cases, whereas floral visitors did so only for one case. Changes in superior competitor’s identity within species pairs are denoted with a blue asterisk.

## Discussion

Until recent years, ecologists have considered that competition mediated by soil conditions has greater potential than aboveground multitrophic interactions to promote or impede plant diversity maintenance. Although this view has been strongly challenged by theoretical and review efforts (Chase *et al.* 2002; Chesson & Kuang 2008), lack of direct empirical evidences has limited the awareness that above and belowground drivers of plant competition can produce symmetrical effects on diversity. Our ability to combine recent advances in coexistence theory with plant population models and detailed observations of spatial variation in soil salinity content and floral visitor frequency during the species’ growing season provide direct evidences that both drivers can equally modify the likelihood of plant coexistence. We particularly observed in our study opposing effects on plant coexistence via direct changes in per capita seed production and indirect changes in competitive responses. While floral visitors promoted plant diversity at the neighbourhood scale, soil salinity drove competitive dominance. Nonetheless, the identity of the superior competitor changed when soil salinity effects on plant competition were considered, which indicates how belowground drivers can maintain plant diversity at larger scales by spatial changes in soil conditions.

At the neighbourhood scale, floral visitors consistently promoted species coexistence by reducing the niche differences needed to overcome fitness differences between species pairs (Fig. 4). The positive effect of floral visitors on diversity did not only occur due to a facilitative effect expected from mutualistic interactions. Rather, we observed both positive and negative effects on plant fecundity. For instance, floral visitors strongly increased the seed production in the absence of competition and reduced to a lesser extent the negative effect of competition on the seed production of *C. fuscatum* individuals. At the other extreme, floral visitors reduced the fecundity of species such as *L. marocanum* and *P. paludosa* by both reducing seed production in the absence of competition and increasing their sensitivity to competition (Fig. 3b and 3c). In our study, the distinct floral visitors’ assemblages observed for these species help to explain these different effects. While bee and fly pollinators mainly visited *C. fuscatum* individuals, beetles were the main visitor of *L. marocanum* individuals. Although it has been observed that some beetles can be good pollinators in Mediterranean ecosystems (Bartomeus *et al.* 2008), most of the beetles in our study were pollen feeder species belonging to the genera *Chrysomelidae* and *Melyridae* (Wäckers *et al.* 2007).

Critically, the equalising effect of floral visitors on plant coexistence likely happened because positive and negative effects were influenced by the species’ competing ability. The negative effect of floral visitors occurred for those species that were on average superior competitors, whereas positive effects occurred for the inferior competitors. This process arises from the fact that our system was dominated by non-specialist interactions and may be a common scenario in this type of systems. Beetles acted as herbivores that tend to focus on the most abundant resource, and therefore target the most abundant species (Table 1). Meanwhile, species with high pollination dependence (i.e. self-incompatibile mating system), tend to be subdominant and the ones that benefit substantially from pollinator’s visits (Tur *et al.* 2013). Although we did not observe that floral visitors in our system produce a stabilising effect on plant coexistence (Fig. 4a, none of the arrows move to the right side of the graph indicating increased niche differentiation under the effect of floral visitors), this does not mean that this density-dependent effect can occur in more specialised systems. In fact, recent studies suggest that equalising and stabilising effects occur in combination. Many plant species trade-off between being sufficiently specialised to differentiate in their pollination niche, while being able to attract a sufficient number of mutualistic partners (Vamosi *et al.* 2014; Coux *et al.* 2016).

Conversely to floral visitors, soil salinity promoted competitive exclusion at the neighbourhood scales of species interactions and it did so by reducing niche differences and increasing fitness differences among species pairs (Fig. 4b). Nevertheless, the identity of the competitive winner changed across contrasting soil salinity conditions. For instance, *B. macrocarpa* and *L. maroccanus* were competitive winners under low soil salinity concentrations but they were competitive losers against *P. paludosa* under high soil salinity concentrations. For the particular case of *P. paludosa*, competitive superiority came mostly from the strong positive effect of salinity in reducing its sensitivity to competitive interactions rather than from an increase in the species’ ability to produce seeds in the absence of neighbours (Fig. 3f). The consistent effect of soil salinity in determining competitive exclusion across species pairs predicts reduction of species diversity in homogeneous landscapes under constant soil salinity conditions, favouring species that either prefer or refuse salt. But in heterogeneous landscapes like our system, diversity is maintained because of the species’ inability to be competitive superiors across all soil salinity conditions. Indeed, these results align with the well-known effect of environmental heterogeneity on promoting diversity (Chesson 2000), and agree also with spatial patterns of species turnover found for very similar salty grasslands in other Mediterranean areas (Pavoine *et al.* 2011). Yet, our results highlight that competitive interaction rather than niche partition (see Rosenzweig 1995; Allouche *et al.* 2012) is likely the main mechanisms driving documented patterns of species turnover.

Our methodological approach is novel in showing how to incorporate the effect of different abiotic and biotic variables into the estimation of niche and fitness differences between species pairs from models that describes species population dynamics via species’ vital rates and interactions coefficients. Our methodology is readily available to be extended to consider other kinds of interactions beyond the scope of this study such as herbivores, leaf pathogens and roots mutualisms. These interactions could potentially explain changes in fecundity of those species for which soil salinity and floral visitors did not have a significant effect (Landwehr *et al.* 2002; Mitchell 2003; Pan *et al.* 2015). More generally, future research needs to fully consider the influence of multiple abiotic and biotic aboveground and belowground interactions on niche and fitness differences for significantly advancing our fundamental knowledge of how diversity is maintained.

Another important step when studying the effect of multitrophic interactions on plant coexistence is to move from direct pairwise effects to include “higher order effects” among species (Mayfield & Stouffer 2017). Higher order efffects occur when the presence of a third species changes per capita competitive interactions within a species pair. One main challenge to this is to achieve sufficient sampling size to capture the variability in species composition and multitrophic interactions (Levine *et al.* 2017). For example, our study does not support such complexity view as model selection by AIC highlighted a common effect of floral visitors and soil salinity across species. However, this could be caused because we measured interactions in a relatively dry year and the abundance of some floral visitors groups such as bees and butterflies were low. This last point also makes us aware that climatic variability across years is another layer of complexity that we do not include in our study. Variation between years in the amount of rainfall can change the spatial configuration of soil salinity conditions as well as change the strength and the specificity of the effect of floral visitors on competitive interactions.

In summary, our study shows that soil conditions and multitrophic interactions represented by floral visitors have contrasting outcomes in determining coexistence at the neighbourhood scale of plant species interactions. While soil salinity promotes competitive exclusion, floral visitors promote coexistence. These differences were mostly explained by equalising processes rather than by stabilising processes. Nevertheless, soil salinity promotes coexistence over larger scales by changing the identity of the competitive winner under contrasting soil salinity conditions. Together, our results highlight that abiotic and biotic determinants of plant diversity are operating at distinct scales. Future research addressing more complexity by including several abiotic and biotic factors, high order effects, and climate variation are needed.

## Author’s contribution

I.B. and O.G. designed the study. J.B.L and O.G. conducted fieldwork. All authors analysed the results and wrote the manuscript.

## Acknowledgments

O.G. acknowledges postdoctoral financial support provided by the European Union Horizon 2020 research and innovation program under the Marie Sklodowska-Curie grant agreement No 661118-BioFUNC. I.B and O.G. acknowledge also financial support provided by the Spanish Ministry of Economy and Competitiveness, Explora Program (CGL2014-61590-EXP, LINCX). J.B.L thanks Francisco Rodriguez-Sanchez for his useful tips and templates to develop all this work in markdown. We thank Curro Molina and Oscar Aguado for their help in field work and identifying insect species. We also thank the Doñana NP staff for granting access to Caracoles Ranch, and Manu Saunders for her manuscript review.

## Appendices

**Apendix S1.**
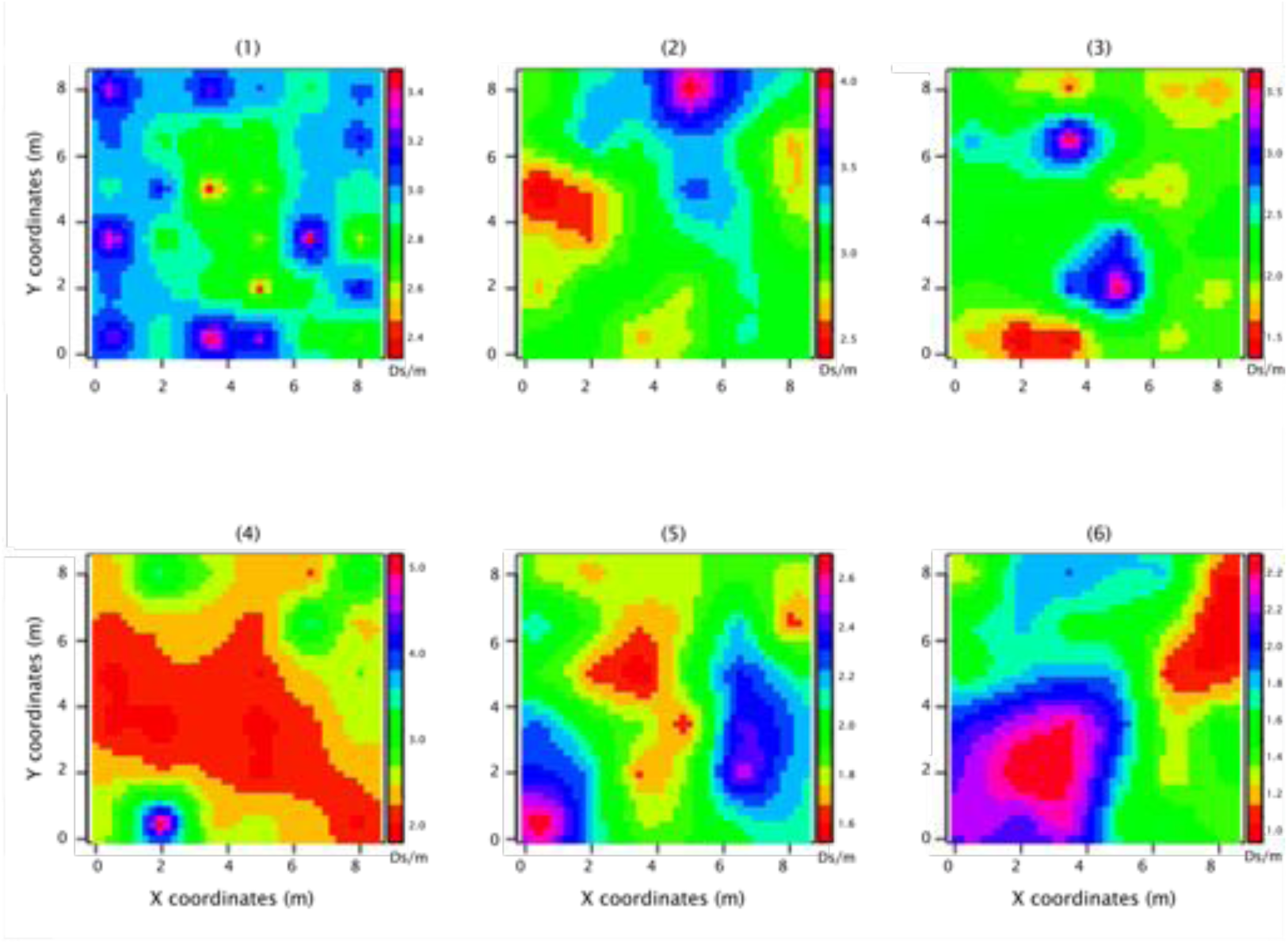
Spatial variation of soil salinity at the plot scale for six of the nine plots included in the experiment. Note that there is high variation in soil salinity content within plot and the spatial configuration changes across plots.

**Apendix S2.**
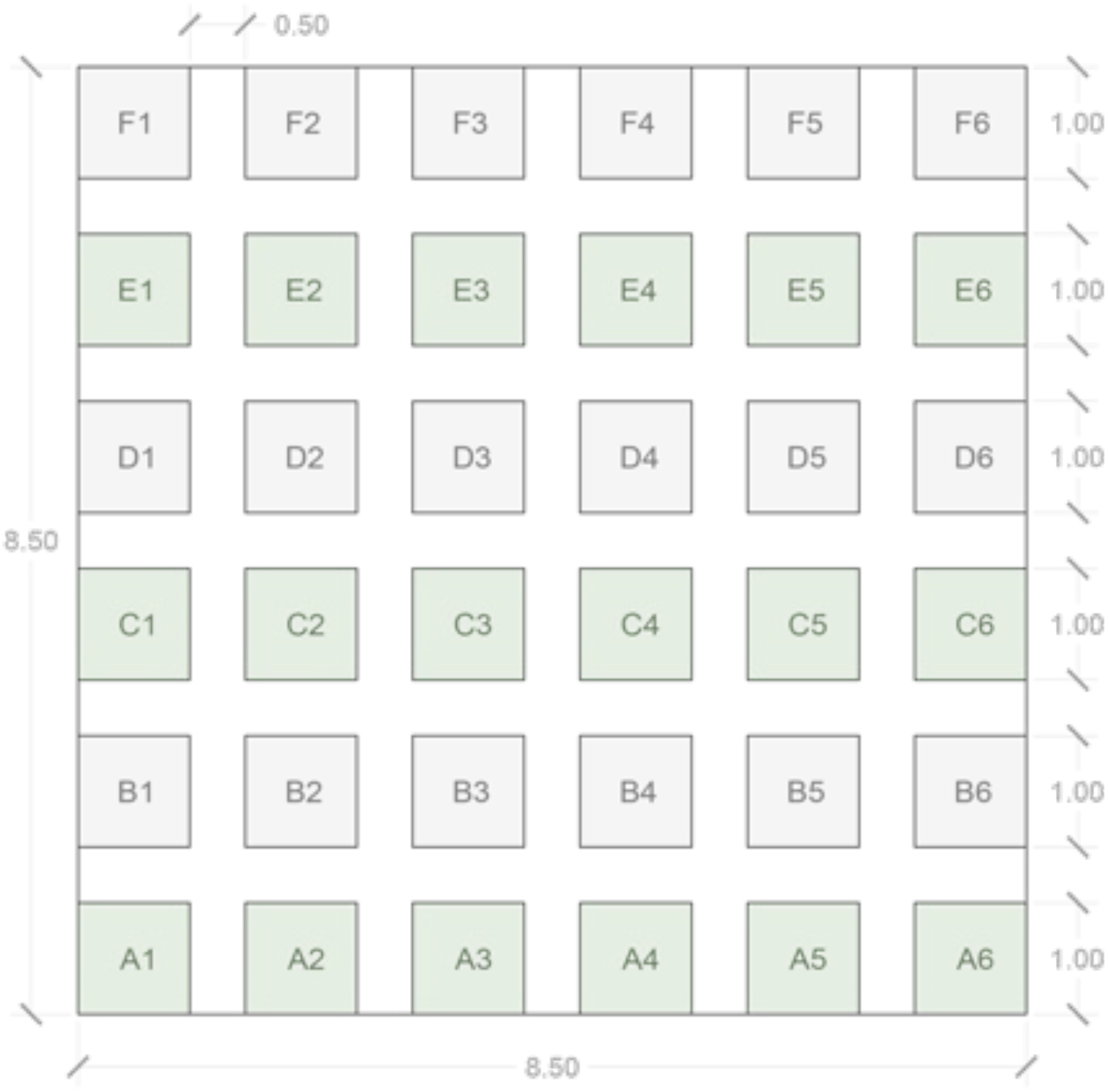
Design of the nine plots included for the study. Each plot of 8.5 × 8.5 m was divided into subplot of 1 × 1 m. Within these subplots, we measured competitive interactions and the effect of salinity and floral visitors of species’ fecundity. Between subplots we left aisles of 0.5m for moving around the plot without producing any negative effect at the subplot vegetation.

**Apendix S3.**
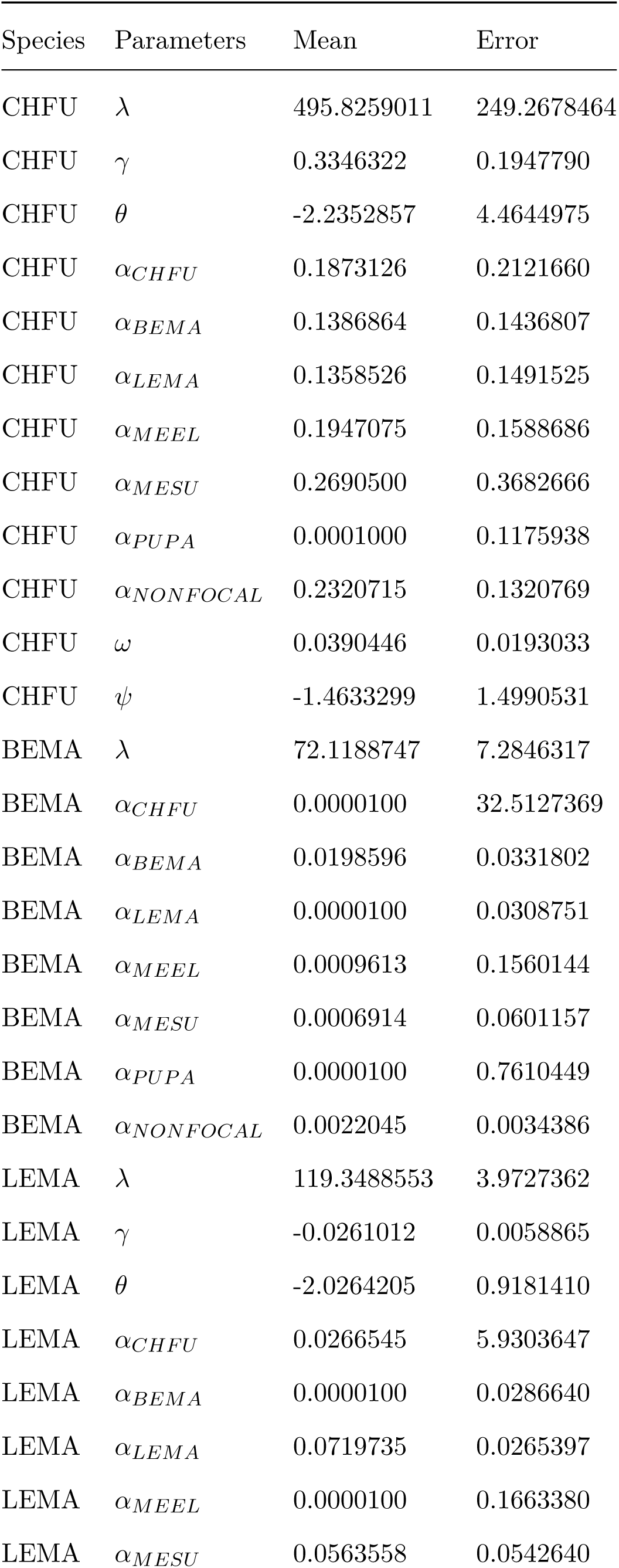
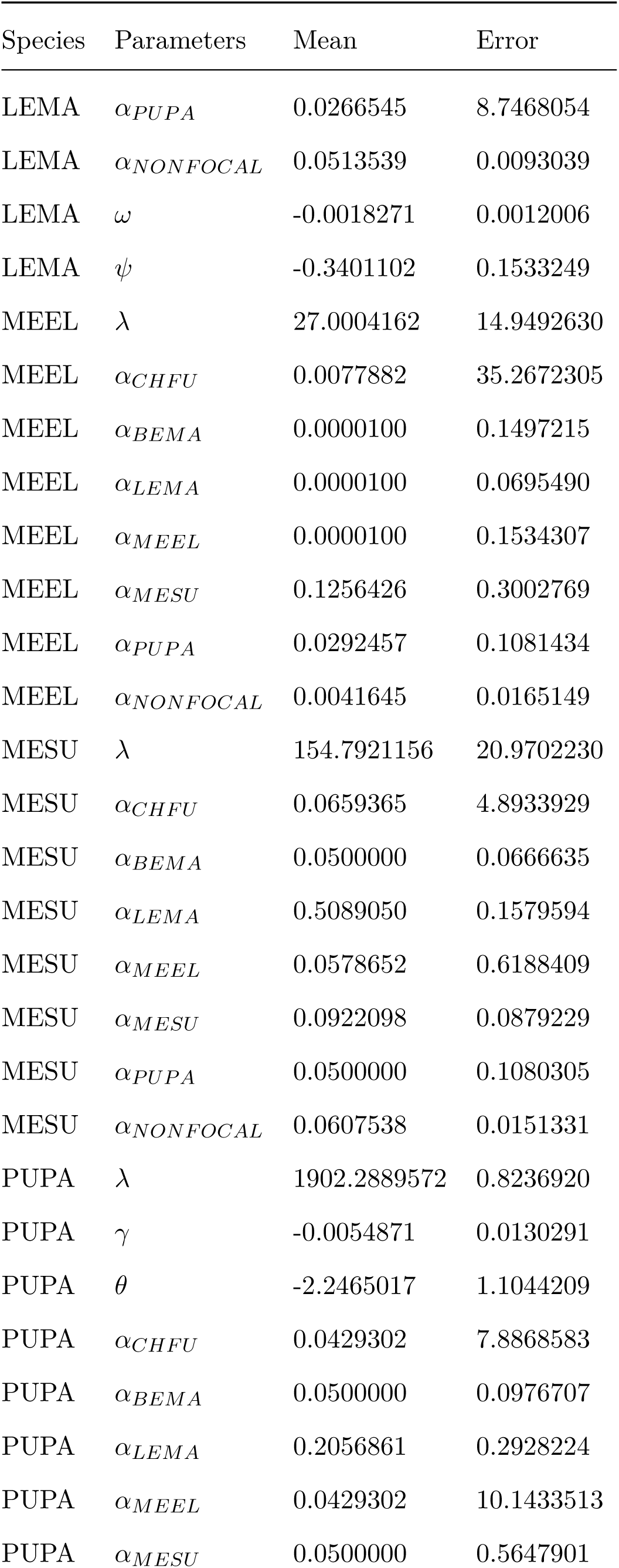
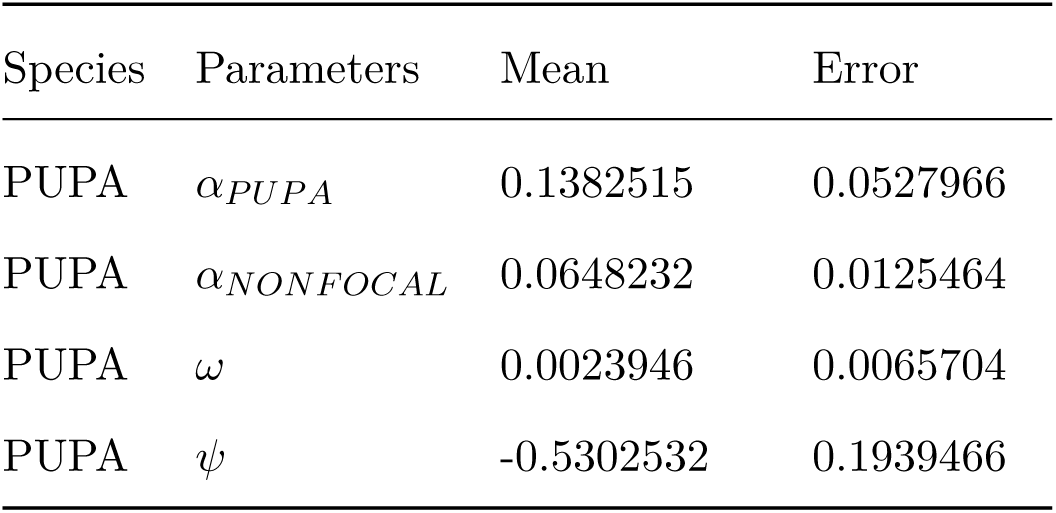
Mean and standard error of each parameter for the best model selected with AIC.

**Apendix S4.**
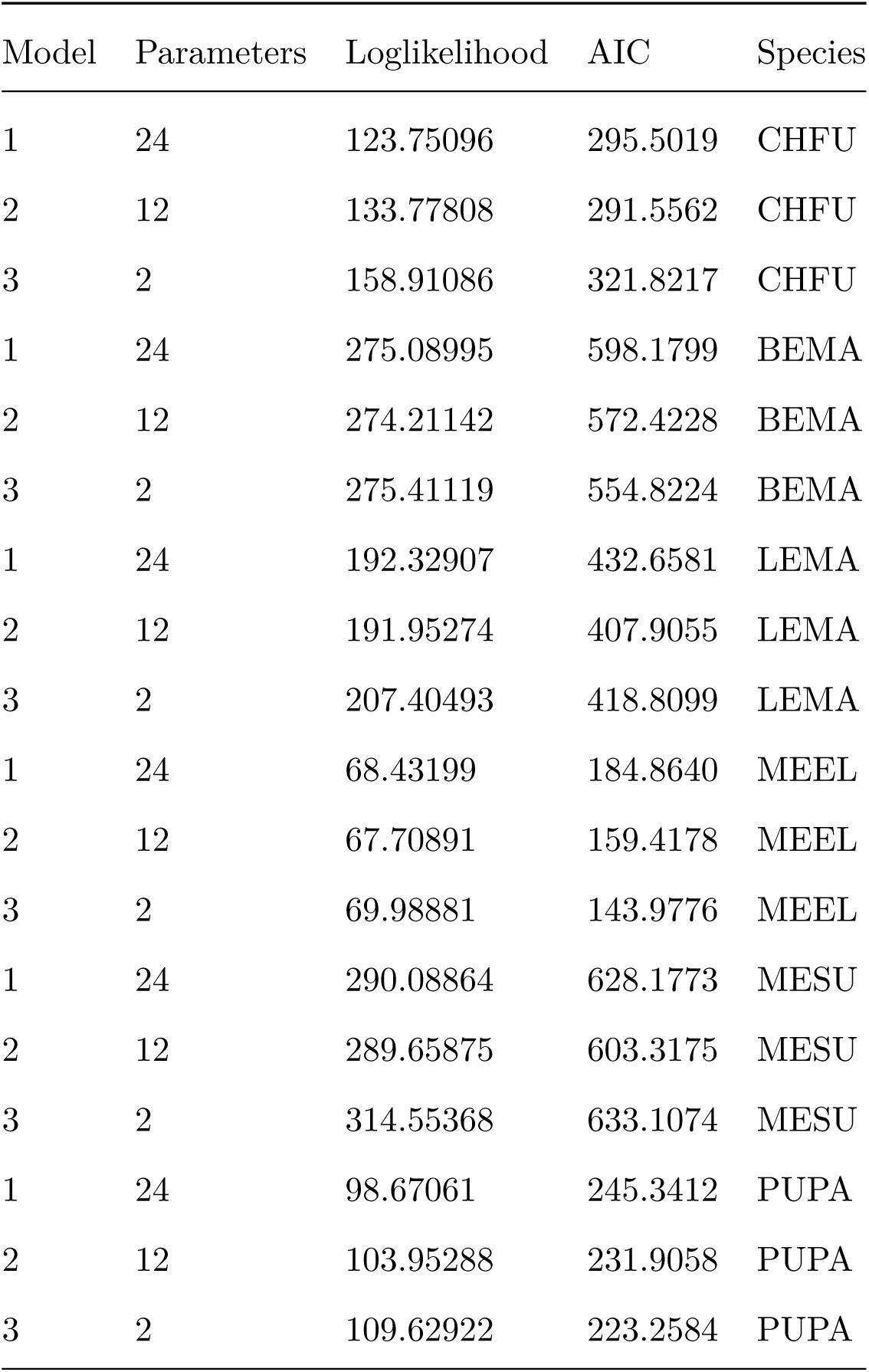
AIC (Akaike Information Criteria) values for the three models considered for each focal species. Species codes are provided in Table 1.

